# Temporal Multiple Kernel Learning (tMKL) model for predicting resting state FC via characterizing fMRI connectivity dynamics

**DOI:** 10.1101/367276

**Authors:** Sriniwas Govinda Surampudi, Joyneel Misra, Gustavo Deco, Raju Bapi Surampudi, Avinash Sharma, Dipanjan Roy

## Abstract

Over the last decade there has been growing interest in understanding the brain activity in the absence of any task or stimulus captured by the resting-state functional magnetic resonance imaging (rsfMRI). These resting state patterns are not static, but exhibit complex spatio-temporal dynamics. In the recent years substantial effort has been put to characterize different FC configurations while brain states makes transitions over time. The dynamics governing this transitions and their relationship with stationary functional connectivity remains elusive. Over the last years a multitude of methods has been proposed to discover and characterize FC dynamics and one of the most accepted method is sliding window approach. Moreover, as these FC configurations are observed to be cyclically repeating in time there was further motivation to use of a generic clustering scheme to identify latent states of dynamics. We discover the underlying lower-dimensional manifold of the temporal structure which is further parameterized as a set of local density distributions, or latent transient states. We propose an innovative method that learns parameters specific to these latent states using a graph-theoretic model (temporal Multiple Kernel Learning, tMKL) and finally predicts the grand average functional connectivity (FC) of the unseen subjects by leveraging a state transition Markov model. tMKL thus learns a mapping between the underlying anatomical network and the temporal structure. Training and testing were done using the rs-fMRI data of 46 healthy participants and the results establish the viability of the proposed solution. Parameters of the model are learned via state-specific optimization formulations and yet the model performs at par or better than state-of-the-art models for predicting the grand average FC. Moreover, the model shows sensitivity towards subject-specific anatomy. The proposed model performs significantly better than the established models of predicting resting state functional connectivity based on whole-brain dynamic mean-field model, single diffusion kernel model and another version of multiple kernel learning model. In summary, We provide a novel solution that does not make strong assumption about underlying data and is generally applicable to resting or task data to learn subject specific state transitions and successful characterization of SC-dFC-FC relationship through an unifying framework.

## 1. Introduction

Since its discovery over two decades ago, there has been a keen interest in investigating the spontaneous intrinsic activity of the human brain. This activity is measured via slow fluctuations in the functional magnetic resonance images (fMRI) when subjects are at rest and not engaged in any task [1]. These fluctuations are highly correlated and discovery of meaningful large-scale functional networks within these correlations led to the use of resting-state fMRI (rsfMRI) to discover human brain function(s) [2, 3]. The resulting matrix of pairwise correlations between regions of interest (ROIs) is termed the functional connectivity (FC) matrix. Many studies of FC have discovered distinct sets of functionally related regions exhibiting temporal correlation in their activities, commonly known as resting state networks (RSNs) [4, 5, 6, 7].

Diffusion tensor imaging (DTI), complementing fMRI, captures the white matter streamlines that form the anatomical pathways along which neural activity spreads over the brain [8, 9]. The topography of the brain anatomy, called the structural connectivity (SC), is estimated by counting the number of streamlines connecting a pair of ROIs. Over the last decade, understanding the link between anatomical topology and neural activity has been an important question in neuroscience. How the relatively static SC sculpts the FC over the entire scan duration has been a challenging open research problem in the brain connectome research domain. Initial studies provide evidence that the underlying structural topology largely explains the grand-average functional connectivity [10], the missing link being dynamics. Whole brain computational models aid study and simulation of the temporal dynamics over the structure.

Extant whole-brain models advancing our understanding of the SC-FC link can be broadly categorized as follows: (*i*) models incorporating non-linear dynamics [11, 12], (*ii*) graph theoretic models [13, 14, 15], (*iii*) models at the boundary of biophysics and graph-theoretic abstractions [16, 17, 18]. Becker et al. [15] mapped spectral signatures of the structural and functional topologies based on indirect structural walks of the neural activity. Abdelnour et al. [16] proposed a graph-diffusion framework relating linear diffusion equation of the neural activity over the structural topology to random walks of the activity over the structure. Surampudi et al. [17] proposed abstraction of non-linear diffusion equation into combinations of multi-scale diffusion to map a subject’s SC-FC.

Over the last decade, several studies of rsfMRI revealed fluctuating spatial patterns which appear and dissolve with time, highlighting the spatiotemporal repertoire of spontaneous brain activity [19, 20]. Attempts at discovering temporal dynamics of rsfMRI can be broadly categorized in the following terms: (i) dynamic functional connectivity (dFC) studies using sliding window approaches providing sequence of windowed FC (wFC) matrices that in turn identify stable transient patterns of functional connectivity fluctuations, called *latent states*, [21, 22, 23, 24], and (ii) Bayesian approaches applied on the time-series themselves [25, 26, 27] which discover latent states in terms of multivariate Gaussian density distributions of the temporal signals. A general perspective is that the neural activity during a task, although being in a high-dimensional space, follows trajectories in a lower-dimensional task-specific manifold during the functional dynamics [28]. This sufficiently motivates the presence of a lower-dimensional manifold for rsfMRI as well.

Moreover, the question of how a relatively fixed anatomical structure supports the rich spatiotemporal dynamics is still elusive. Abdelnour et al. [18] have extended their graph-diffusion framework for characterizing SC-dFC relationships. However, theoretical models incorporating principled amalgamation of structural topology and dynamics of rsfMRI are essential. Here, we propose an innovative solution for characterizing the SC-dFC-FC relationship. This is achieved by proposing two novel constructs: (*i*) discovery of a lower-dimensional manifold that represents the latent structure of the temporal dynamics, (*ii*) temporal multiple kernel learning (*tMKL*) model that learns the SC-dFC mapping, and (*iii*) generation of latent time series for dFC-FC mapping. The proposed solution estimates grand average FC (gFC or FC) from SC by predicting dFC along with capturing the temporal evolution. Temporal evolution is characterized by using a first-order Markov model between consecutive state transitions. This model is used for generating a long state sequence using the steady state distribution of the Markov random walk. This state sequence is further replaced by sequence of corresponding state-specific FCs generated by the tMKL model. Finally, these state-specific FCs are factorized to recover a latent time-series sequence. gFC is then computed on the reconstructed latent time-series and compared with the empirical FC. The proposed model recovers the FCs that are very close to empirical FCs as the state-specific FCs recovered with the tMKL model enable realization of subject-specific functional dynamics. Further, various perturbation experiments demonstrate the robustness and validity of the proposed scheme. This state sequence is further replaced by sequence of corresponding state-specific FCs generated by the tMKL model. Finally, these state-specific FCs are factorized to recover a latent time-series sequence. gFC is then computed on the reconstructed time series and compared with the empirical FC. The proposed model recovers the gFCs that are very close to empirical FCs as the state-specific FCs recovered with the tMKL model enable realization of subject-specific functional dynamics. Further, various perturbation experiments demonstrate the robustness and validity of the proposed scheme.

The specific contributions of the work are the following:

1. Novel approach for learning the SC-FC mapping through characterizing the dynamic functional connectivity (dFC) over time windows.
2. Proposal of a novel multiple diffusion kernel model that learns to predict state-specific FCs from SC (tMKL model).
3. Estimating the latent fMRI time series by using the Markov transition probability matrix in conjunction with the tMKL model.

The rest of the paper is organized as follows. In the next section we present the details of the proposed solution. In the subsequent sections we present the details of the neuroimaging data set used, qualitative and quantitative evaluation results along with explanation for the choice of model parameters. Finally, we conclude by pointing out limitations and future research directions.

## 2. Materials and methods

### 2.1. Dataset

Resting state fMRI as well as corresponding diffusion weighted imaging (DWI) data were collected at the Berlin Center for Advanced Imaging, Charité University, Berlin, Germany. The dataset consisted of structural connectivity - functional connectivity (SC-FC) pairs of total 46 subjects used in this study. In summary, all the participants underwent resting state functional imaging (no task condition) with eyes closed for 22 minutes, using a 3T Siemens Trim Trio scanner and 12 channel siemens head coil (voxel size 3 × 3 × 3 mm). Each fMRI resting state data amount to a total of 661 whole brain scans (time points recorded at TR=2s)were obtained during the resting state functional magnetic resonance imaging (rs-fMRI) session. Thus the blood oxygen level dependent (BOLD) time-series signal available for each participant has 661 time points aggregated across 68 regions of interest (ROIs) as per the Desikan-Killiany brain atlas [29]. The diffusion weighted tensors (TR=750 ms, voxel size 2.3 × 2.3 × 2.3 mm) computed from the dwMRI data recorded with 64 gradient directions were subjected to probabilistic tractography as implemented in MRTrix [29] in order to obtain subject specific sturctural connectivity (SC) matrices. Masks derived from high-resolution T1-images were used to determine seed-and target-locations for fibers in the grey/white matter-interface (GWI). SC matrices contains connection streamlines obtained based on the fiber tracking algorithm with various assumptions based on known limitations imposed by anatomy, notably the size of the GWI of each region. Further image acquisition, choice of scan parameter details and data pre-processing methodology adopted are all available in [30].

### 2.2. Proposed model

In this section we describe in detail the whole pipeline of the proposed parametric-model to map the relation between SC and FC using resting state fMRI data. The proposed model considers the importance of the underlying anatomical constraints to generate the temporal richness as well as to characterize and assess whole-brain FC dynamics. Figure 1 shows a flowchart of the essential elements of the whole pipeline. Proposed model partitions aspects of the whole-brain dynamics essentially into two parts: characterizing temporal dynamics through identification of latent transient states and linking them to the underlying structural geometry. These two aspects are parameterized using a novel combination of unique methods. The model utilizes wFCs (steps 1. – 2.) for identifying states and from the resultant SC-wFC pairs, the relationship between the structure and functional dynamics is learned (steps 3. – 5.). Once these two parts successfully characterize the above mentioned aspects by tuning respective parameters, the model is tested for its generalizability using unseen test subjects (steps 6. – 9.).

**Figure 1:**
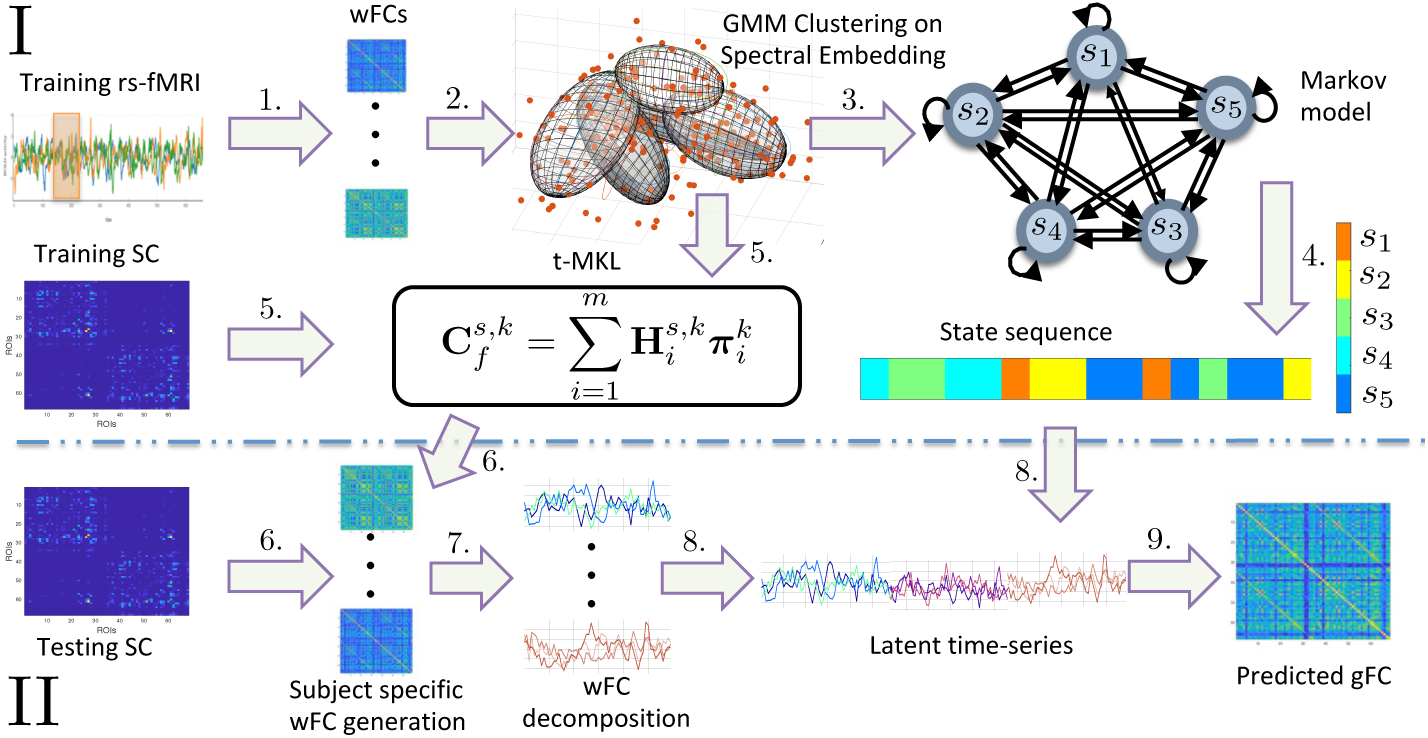
Outline for temporal Multiple Kernel Learning (tMKL) model. Figure shows the entire pipeline for predicting grand FC for a testing subject. The model incorporates subject specificity along with temporal variation characterization. Part (I) of the model, training phase, consists of three blocks. The first one, learns temporal variations in terms of distinct states via GMM clustering over the underlying manifold of wFCs (steps 1. and 2). The second block utilizes the empirical transitions between these distinct states and captures dynamics in the first order Markov chain (steps 3. and 4). The third block learns subject-specificity by modeling each state by its MKL model [17] (step 5.). Part (II) of the model validates its generalizability on unseen subjects. Importantly, only SC of a testing subject is required (step 6). Each state for the testing subject is characterized in step 7. Each state-specific predicted FC is decomposed into a latent time series which are then concatenated using the steady state distribution of the Markov chain (steps 4. and 8). Finally, grand average FC, static FC, is predicted for that subject (step 9).

For identifying latent states within the dynamics, we discover the underlying globally non-linear manifold that spans all the wFCs (step 2*.a*), thus recovering the lower-dimensional space for meaningful characterization. We employ a probabilistic framework for estimating the number of states and the shape of each state in the lower-dimensional space, ensuring soft assignments of wFCs to its neighboring states (step 2.6). These soft assignments are further used to estimate the transition dynamics between these states (step 3. – 4.). With respect to second aspect of the model, we adapt the multiple kernel learning (MKL) framework [17] for parameterizing the dependence of SC on wFCs for each state (step 5.). We observe that the parameters to be learned form a non-convex combination, necessitating an iterative algorithm. Thus we formulate the learning objective into an optimization formulation and adapt an iterative algorithm for solving this non-convex combinations of parameters.

The model predicts state-specific FCs (sFCs) for a test subject (step 6.). These sFCs are decomposed into a latent time-series (step 7.) which is concatenated using the relative frequency of occurrence of states to generate a global time-series for calculating the static FC of a subject (step 8.). Thus, for a new subject, given the SC, static FC along with its state-specific FCs are predicted by the proposed model (step 9.).

In the subsequent subsections we elaborate each part of the proposed model. From now on, let 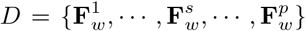 be the set of all wFC matrices obtained by sliding a window of fixed size *ω* over the *n*-dimensional fMRI time-series belonging to all the training subjects.

#### 2.2.1. Spectral Embedding, step 2.a

We propose to soft-cluster these wFC matrices into *K* states, first by vectorizing the lower triangular part of a wFC matrix into a column vector of size 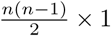. These wFCs may be sparsely spaced in a higher-dimensional space, but might originally lie on an intrinsic globally non-linear manifold [31]. Spectral embedding method is employed to reduce the dimensionality of the data, by finding a mapping to a lower dimensional manifold over which these wFCs reside [32]. The graph constructed over the vectorized wFCs provides a discrete approximation of the continuous manifold. The solution embedding is provided from the eigenmaps (eigenvectors) of the Laplacian operator over the graph, which approximates a natural mapping onto the entire manifold. The Laplacian eigenmaps preserve the local structure in the graph, thus keeping the solution embedding robust to outliers and noise.

The spectral embedding method is applied as follows. Firstly, an affinity matrix is created by applying a radial basis function over the L1 distance between every pair of wFCs. This matrix captures pairwise relationship between wFCs in a relational graph. Next, we form the corresponding normalized graph Laplacian matrix and use the eigenvectors corresponding to its lowest *K* eigenvalues to define the basis vectors of embedding space [33, 34, 35]. The value of these eigenvectors against each wFC represent its resulting transformation into the embedding space. Finally these *K*-dimensional embedded wFCs are clustered using Gaussian Mixture Model (GMM), as explained in the next subsection.

#### 2.2.2. GMM Clustering, step 2.b

Following the discovery of an approximation to the continuous lower-dimensional manifold, we now parameterize the local density distribution of wFCs over the manifold using a probabilistic framework, Gaussian mixture model (GMM) [36]. Gaussian mixture model is a factor analysis model that represents the probability density of a sample as a weighted combination of component Gaussians. Such a representation facilitates GMM to represent a large class of sample distributions. Specifically, distribution of wFCs over the manifold are modeled as a GMM.

Let the density of 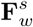 be a linear combination of *K* component Gaussian densities, represented as follows:

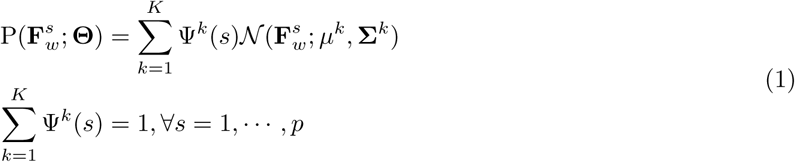

where P denotes the probability density of a wFC. Each component Gaussian is a K-variate Gaussian probability density function of the form:

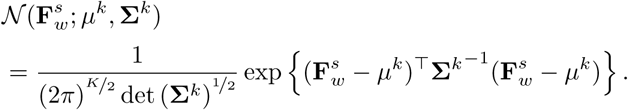

GMM thus represents the manifold as a set of Gaussian densities and parameterizes it in terms of Θ:

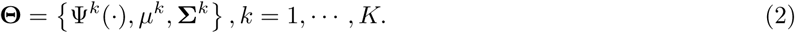

As the collection of these component Gaussians forms the manifold, the component Gaussians can be interpreted as a *latent transient state* visited by the brain. Each state is a Gaussian but at different locations and with different shapes governed by *μ^k^* and Σ*^k^*, respectively in the manifold.

#### 2.2.3. State Transition Markov Model, step 3

As described in the previous section, the wFCs are quantized into finite states *S* = {*s*_1_, …·*s_K_*} by GMM clustering. Each wFC sequence now corresponds to a cluster-label (state) sequence and transitions between these states is representative of the dynamics in the BOLD rsfMRI time series. We assume first-order dependence among these transitions and learn the Markov transition probability matrix, **T***_K×K_* by estimating the state transitions from the training data.

Figure 2 shows a depiction of Markov model for *K* = 5 and the corresponding transition probability matrix. Each edge *t_i,j_* captures the probability of transition from state *i* to state *j*. Similarly, self-loop edges *t_i,i_* depict the probability of remaining in the same state. For each state *i* we compute *t_i,j_* by counting the number of first-order transitions to state *j* in the state sequence. Finally, we normalize each row of **T** to make it a valid transition probability matrix. In the testing phase, the Markov matrix learned on training wFCs is used to generate a random state sequence, to eventually construct the latent time-series for testing subjects. As any Markov chain converges to its steady state distribution with time regardless of its initial distribution, we find the steady state distribution over the transition matrix and use this distribution as frequency of occurrence of states over the time course. This gives us a state transition sequence for a test subject. Along with the state transition model which captures the dynamics of the latent states, a model that relates anatomical structure to these states is required. In the next section, we propose a temporal multiple kernel learning (tMKL) model that learns this mapping.

**Figure 2:**
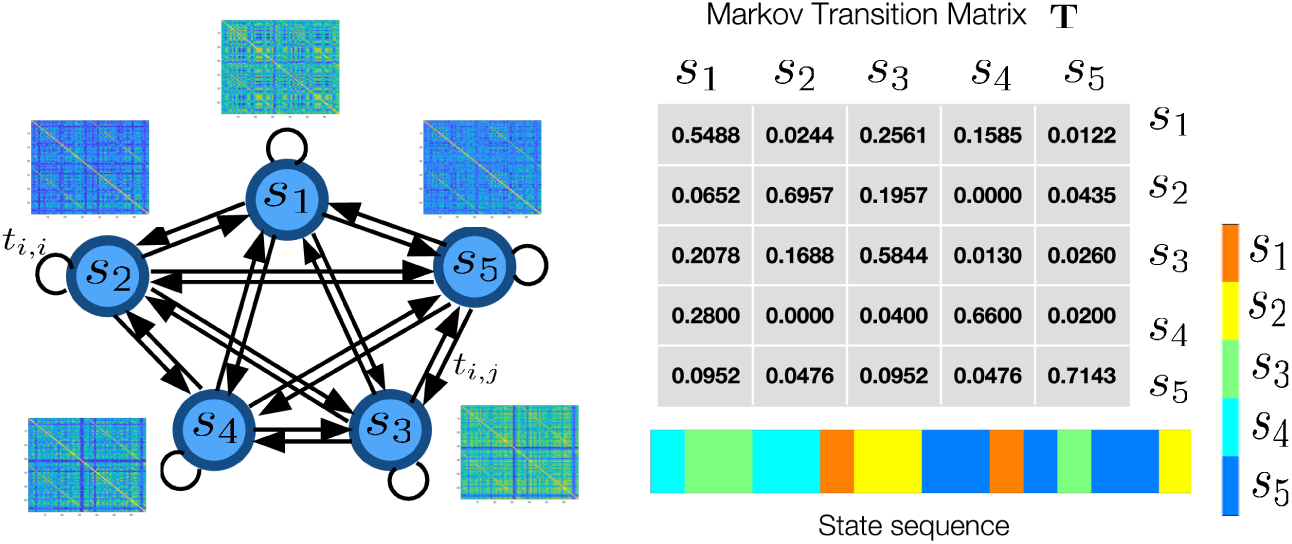
Graphical depiction of proposed Markov state transition model. An illustration of the first-order Markov chain used as a part of the proposed tMKL model. Each state has its unique distribution of FCs, represented as a Gaussian in the embedding space, from which subject-specific FCs can be sampled. The corresponding transition matrix (for K=5) and an example state sequence generated with a Markov random walk over the transition matrix is also depicted.

#### 2.2.4. tMKL Model, step 5

Mean regional activities of all regions are assumed to be in a random walk over the SC graph. This phenomenon is represented by a linear differential equation whose analytical solution is the diffusion kernel over the graph defined by SC which is hypothesized to be representing FC [16]. [14] discovered that physical diffusion over such large scale graphs exhibits multi-scale relationships with FC, thus a linear combination of multiple diffusion kernels is considered more representative of FC (this model is referred to as MKL_NIPS from now on). The linear combination coefficients are scalar values which equally weigh all regional activities at each diffusion-scale. But it may so happen that activities of non-physically connected regions may be modulated by other regions. To represent this phenomenon we introduce the variables **π***_i_*’s of size *n × n*, that capture the inter-regional co-activation patterns at diffusion-scale *i*, ∀*i* =1,…, *m*, *m* being the number of diffusion-scales [17].

Let a diffusion kernel defined at scale *i* be denoted by **H***_i_*.

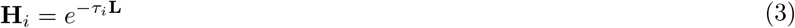

Here, *τ_i_* is the spatio-temporal scale of heat diffusion and **L** is the Laplacian matrix corresponding to the SC. We propose that a wFC matrix can be decomposed into a set of diffusion kernels multiplied with their co-activation pattern:

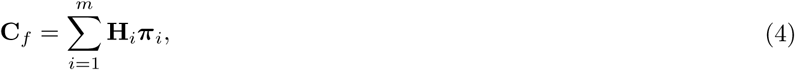

Here, **C***_f_* denotes predicted wFC. We hypothesize that co-activation patterns are distinctly different for each state and hence we add a superscript index *k* (*k* = 1 … *K*) to obtain 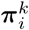. As the parameters 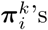 are state dependent, state-specific predicted functional connectivity, 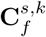, will be as follows:

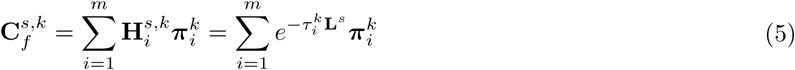

Here **L***^s^* is the Laplacian matrix of the SC corresponding to *wFC*^s^. This results in the following optimization problem for **Π***^k^* and **τ***^k^*:

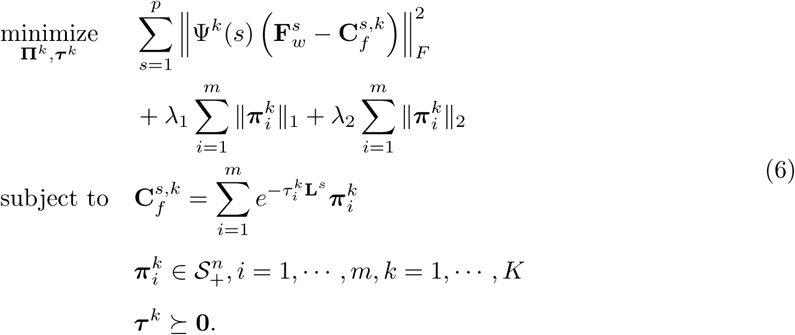

Here, 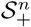 is the convex set of positive semi-definite matrices. The objective function takes the form well known in regression analysis as *least absolute shrinkage and selection operator* (LASSO) that performs both variable selection and regularization. We arrived at the model parameters experimentally, for example, the number of scales *m* is empirically chosen (see Subsection 3.2).

Finally, the model consists of *m* distinct 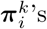 which are learned for each of the *K* states.

#### 2.2.5. Generation of latent time-series for testing subjects, steps 4., 6. – 9

As described in the previous section, we predict the state-specific FC matrix for each of the states using the input SC matrix of the testing subject and the learned tMKL model (step 6.). Based on the learned Markov chain state transition matrix, a sequence of states is generated using the steady state distribution of the transition matrix (step 4.). Each of the state-specific FCs in the resulting sequence is factorized into state-specific latent time-series and concatenate to obtain the latent time-series for the testing subject.

In the training phase, wFCs are obtained by computing Pearson correlation coefficients of the windowed BOLD rsfMRI time-series over various regions. We know that Pearson correlation between two time-series *A, B* is 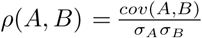. Hence the state-specific wFC matrix works out to be the covariance of its state-specific latent times-series 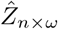. Thus we can factorize a state-specific wFC as follows:

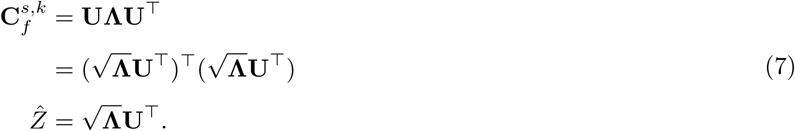

Thus, using Eq. 7, we recover latent time-series matrix 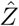 that can be taken as approximated time-series used for obtaining wFC (step 7.). For a testing subject, each cluster-specific wFC is decomposed into latent time-series and these are concatenated into a grand time-series (step 8.). The latent time series are concatenated by considering the steady state distribution of the Markov chain. Steady state distribution is the probability of being in a state which remains the same throughout transitions. Every random walk over the transition matrix approximates this distribution after infinitely long time. Finally, as Pearson correlation is order-agnostic, calculating Pearson correlation matrix of the grand time-series generates the predicted grand-average FC (gFC) for the testing subject (step 9.).

## 3. Experiments & Results

Performance of the proposed model was evaluated in the following setup. A randomly chosen set of half of the cohort (23 participants) was used for training and the other half (23 participants) for testing. We used Pearson correlation coefficient between empirical and predicted functional connectivities (FC) as the measure of model performance in order to keep the measure of model performance consistent with the extant literature. We first compare the performance of the proposed model against several extant methods that provide SC-FC mapping followed by explaining the rationale behind the choice of optimal model parameters. We also conduct k-fold cross validation results and perturbation experiments, the results of which support generalizability of our model to other data splits. The proposed model predicts state-specific FCs which are thereby used to product the gFC. The quality of the gFC prediction is highly dependent upon the reproducibility of states and their transition patterns across multiple train-test splits. Obtaining different set of states in different splits shall attest the robustness of the proposed model at question. Finally, we analyze the states discovered from our model by observing the state-specificity property of the model and compare it with the states learned using k-means algorithm in Allen et al. [21].

### 3.1. Grand average FC (gFC) prediction

We compare the performance of the proposed model with several existing approaches: single diffusion kernel (SDK) model [16], the non-linear dynamic mean field (DMF) model [12] and multiple kernel learning (MKL) model [17]. To our knowledge, ours is the only model that incorporates structural information along with temporal dynamics for predicting grand average FC. DMF and SDK models do not incorporate learning in their formulation and tune the parameters for each subject separately. DMF model inherently captures non-stationarity, therefore it is directly used for gFC prediction without computing wFCs. We estimated the optimal parameters of the DMF and SDK models from the training wFCs and predicted the gFCs of testing subjects using these optimal parameters. The mode of the performance distribution histogram for the training set was used to select the optimal model parameters. Figure 3 shows that tMKL has superior performance compared to the others.

To validate the generalizability of the tMKL model over unseen testing data, we performed k-fold cross-validation experiment whose results are listed in Table 1. These results suggest that performance of our solution is consistent across various splits, hence supporting our claim of generalizability of our model on unseen data.

**Table 1:** Cross-validation experiments suggesting generalizability of tMKL model. Mean *k*-fold cross-validation results for *k =* 2, 3, 5 are shown in the corresponding rows for *k*-values. As the number of training samples increases with the number of folds, the mean performance also increases suggesting that the model is learning well with increased samples ans is able to replicate the same for testing subjects.

| k | fold-1 | fold-2 | fold-3 | fold-4 | fold-5 | mean |
| --- | --- | --- | --- | --- | --- | --- |
| 2 | 0.757 | 0.732 | - | - | - | 0.745 |
| 3 | 0.771 | 0.811 | 0.778 | - | - | 0.787 |
| 5 | 0.785 | 0.809 | 0.813 | 0.809 | 0.808 | 0.805 |

Now, the choice of various model parameters is explained in the next subsection.

### 3.2. Parameter Selection

1. **Choice of size of sliding-window**, *ω*: Within the extant literature, the choice of a suitable sliding window size is an open problem with respect to the analysis of temporal dynamics in rs-fMRI [20]. The sliding window size should be small enough so as not to miss the state transitions and should be large enough to capture the state transitions reliably. Keeping this in mind, we followed Allen et al. [21] by using a sliding window of *ω* = 22 TRs. The window was tapered at the ends by convolving it with a Gaussian of *σ* = 3 TRs and was slid with a stride of 5 TRs to create wFCs.
2. **Choice of GMM parameters:** Each *latent transient state* in which the wFCs lie is represented using a component Gaussian of the GMM. In order to choose the optimal number of these states, *K*, we selected the GMM model corresponding to a minimum BIC score. Bayesian information criterion (BIC) is a statistical measure based on the log-likelihood function used for selecting a model amongst a finite set of alternatives, where the model corresponding to the lowest BIC score is chosen. The plot in Figure 4 shows BIC scores for the models obtained by fitting GMM for a large range of *K* (2 to 19), where the minimum value was obtained for *K* = 12. For each *K*, we ran GMM 100 times and noted the minimum BIC score, these BIC scores were used in the figure. To retain generality of the component Gaussians, we ran our experiments by considering a unique full covariance matrix for each component Gaussian.
3. **Choice of number of diffusion scales for tMKL**, *m*: The scale values were sorted in ascending order, where lower values correspond to local diffusion phenomenon and higher values correspond to global diffusion phenomenon. Scale values lying in-between correspond to intermediate diffusion phenomena. We ran several experiments by varying *m* at powers of 2 from 2 to 32. While the performance for all the scales was reasonable, however in order to carry out comparative analysis with the MKL model, we chose the number of scales as *m* = 16 for all the experiments.

**Figure 3:**
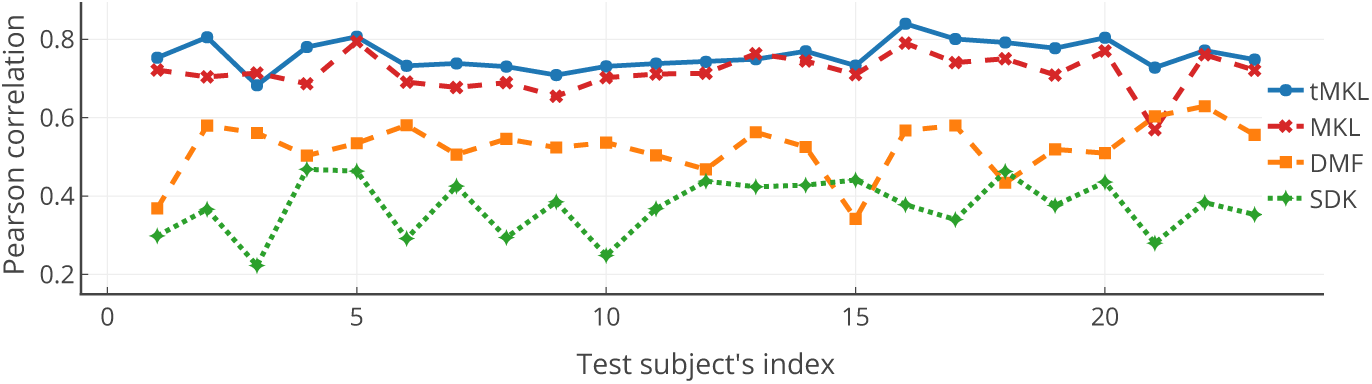
Model performance comparison between tMKL and existing models. Pearson correlation between the empirical and predicted gFCs for all the testing subjects is shown for all models. As can be seen, MKL model outperforms other two models, and tMKL model is at par or better than MKL for all but one testing subjects. Even though there is marginal gain in the overall prediction quality, tMKL provides rich insights into the temporal dynamics thus gaining its superiority over extant models.

**Figure 4:**
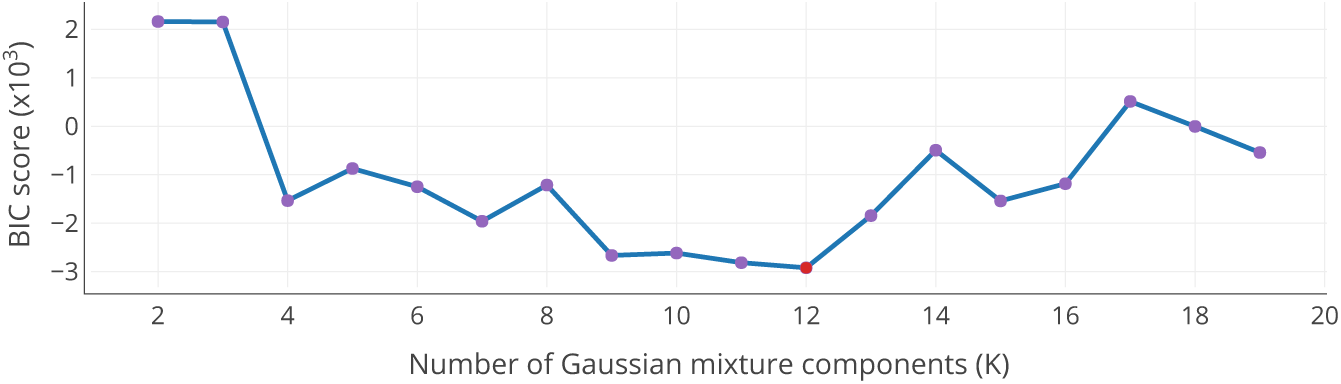
Bayesian information criterion (BIC) score for selecting the number of components in Gaussian mixture model. The GMM is fit over the training wFCs lying in the lower dimensional manifold. The BIC score is reported by varying the number of component Gaussians (*K*) from 2 till 19. The Gaussian mixture model corresponding to *K =* 12 (shown in red) has the lowest BIC score among others and is therefore preferred. The plot shows local minima at *K* = 4, 7, and 15 which may mislead the user while selecting the optimum model. This local minima suggests the choice of number of components in Allen et al. [21].

### 3.3. Robustness of the model

In order to validate the robustness of our model we performed various experiments to assess whether our solution overfits the training data and also whether the prediction of the grand average FC is agnostic to the particular SC matrix.

1. **Reproducibility of states:** As mentioned in Section 2.2.2, GMM yields K soft assignment vectors for the training wFCs. We validated reproducibility of this clustering by ensuring replication of the same for wFCs of the testing subjects. We generated wFCs for all the testing subjects using the sliding window approach. Soft assignment vectors were generated for these testing wFCs using the GMM employed on the training data, which is then used to compute the Markov transition matrix and the corresponding steady state distribution. Figure 5 shows an example of the steady state distribution for our proposed method. We evaluated the similarity between the Markov transition matrix and steady state distributions of the training and testing wFCs by finding the Pearson correlation coefficients. Table 2 shows that the states are highly replicable for multiple train-test splits of the data.
2. **Perturbation experiments:** Each testing subject SC was perturbed *N* = 150 times and using the learned model we predict the grand average FC. We perturbed every SC by randomly generating it from the power law distribution followed by its elements. The generated state-specific wFCs may have non-positive eigenvalues. Here we considered only the real part of the generated time series in order to estimate (predict) grand average FC. Figure 6 shows this observation over all the 23 testing subjects. Box plots for each subject depict the range of correlation values for random SCs. Here we observe less correlations between empirical and predicted gFCs using the perturbed SC, validating that our model respects the topology of input SC. This suggests that the model is not overfitting the data and is sensitive to perturbation in SC.

**Figure 5:**
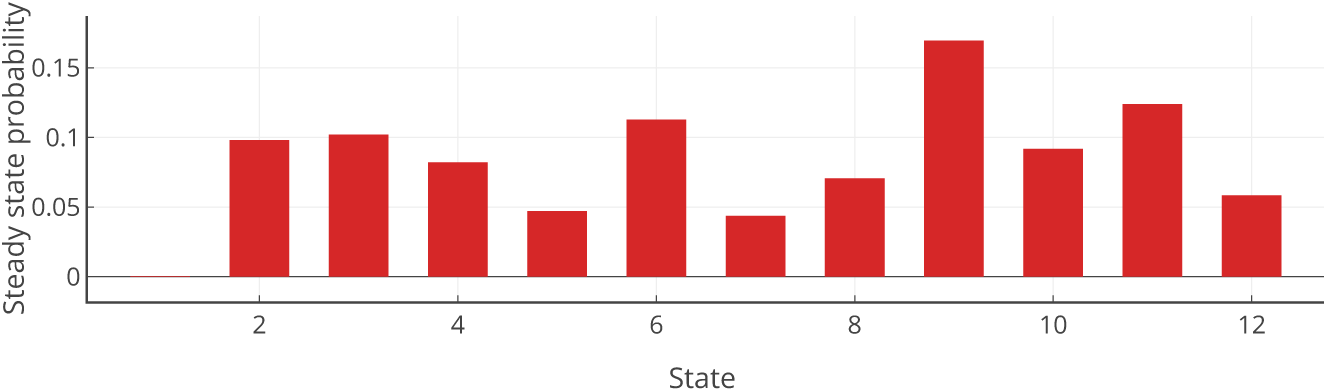
Markov chain steady state distribution. After the states are retrieved using GMM, the Markov chain transition matrix was learned over the resulting state-sequences of the wFCs of the training subjects. The figure shows the steady state distribution of the transition matrix, which represents the probability distribution of occurrence of a state after infinite amount of time.

### 3.4. State-specificity of the tMKL model

**Table 2:** Comparison of Markov chain transition matrix (TM) and its steady state distribution (SSD) between training and testing subjects. Comparison is done computing the Pearson correlation coefficient (*ρ*) and the L2 distance (*e*) between the training-TM, testing-TM and training-SSD, testing-SSD respectively. This experiment is repeated for 11 train-test splits of the data. Consistent high values of *ρ* and low values of e across multiple splits show similarity of the states and their transition behavior across train-test subjects, therefore establishing the reproducibility of states.

| Run Index | $\rho_{\text{TM}}$ | $\rho_{\text{SSD}}$ | $e_{\text{TM}}$ | $e_{\text{SSD}}$ |
| --- | --- | --- | --- | --- |
| 1 | 0.9947 | 0.9509 | 0.1337 | 0.0564 |
| 2 | 0.8683 | 0.8546 | 0.7379 | 0.2703 |
| 3 | 0.9440 | 0.8839 | 0.5120 | 0.1433 |
| 4 | 0.9035 | 0.9809 | 0.7154 | 0.1004 |
| 5 | 0.8624 | 0.9604 | 0.7094 | 0.1332 |
| 6 | 0.9665 | 0.8337 | 0.3824 | 0.1119 |
| 7 | 0.9131 | 0.8563 | 0.6263 | 0.1107 |
| 8 | 0.9746 | 0.6824 | 0.3381 | 0.1521 |
| 9 | 0.9275 | 0.8691 | 0.6671 | 0.0950 |
| 10 | 0.8623 | 0.9599 | 0.7299 | 0.1608 |
| 11 | 0.9777 | 0.9596 | 0.3250 | 0.0482 |
| mean | 0.9301 | 0.8692 | 0.5068 | 0.1358 |
| stdev | 0.0501 | 0.1155 | 0.2093 | 0.0636 |

**Figure 6:**
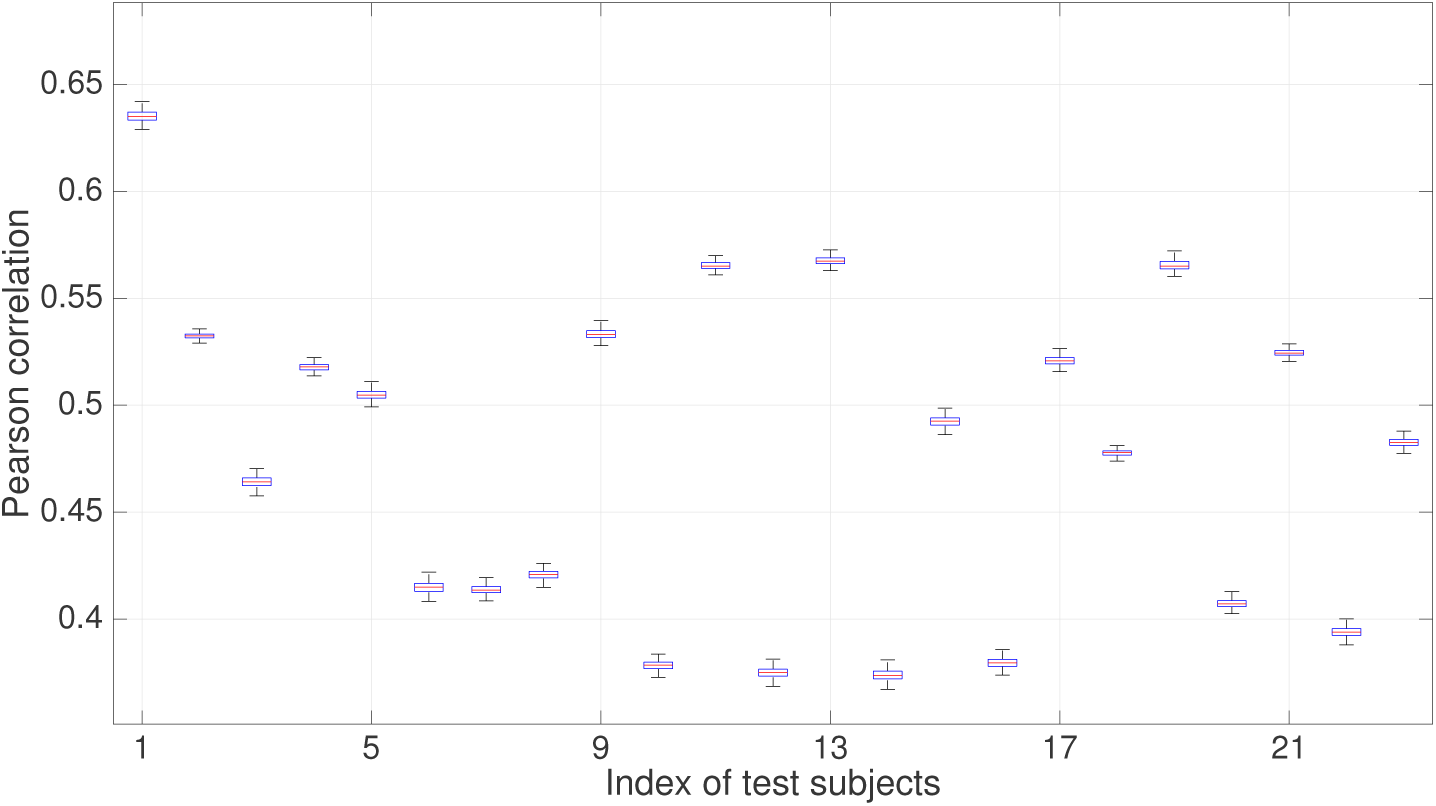
Effect on performance of tMKL model due to perturbation of SC matrices of test subjects. Shown here are the box plots (blue) of Pearson correlation between empirical and predicted grand FCs when SCs are perturbed for all the testing subjects.

In the previous section, we successfully investigated whether the estimated state transition matrix is general enough in the sense of being reproducible with several train-test splits of the data (refer Table. 2). Other critical questions are whether different model components such as the **Π***^k^*’s as well as the predicted FCs are distinct for different states or not. If they are not distinct, the resulting MKL models for different states become redundant. In order to verify the state-specificity of the model, we performed three simulation experiments: i) to show that the learned state-specific model parameters on the training data are distinct for different states, ii) to show that the predicted state-specific FCs during testing phase are also distinct from one another and, iii) to evaluate the accuracy of state-specific assignments of the model prediction using precision and recall measures.

As summarized in Section 2.2.4, the full model consists of estimating *m* =16 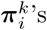 for all the *K* = 12 states (*i* ranging from 1 to 16 and *k* ranging from 1 to 12). We perform a comparison experiment to see whether, for a fixed *i*, 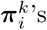 are dissimilar from each other. The results of the first experiment are depicted as *m* =16 similarity matrices in Figure 7. It appears that the learned 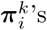 are indeed different for different states, especially in the similarity matrices for global scales (see the top row of Figure 7).

In the second experiment, we verified whether the predicted state-specific FCs during the testing phase are distinct from one another. For a test subject, there are *K* state-specific FCs predicted based on the SC of the subject (see step 6. of Figure 1). In the previous experiment, as the **Π***^k^*’s have been demonstrated to be distinct from each other, given a fixed test SC, the state-specific predictions are also expected to be distinct. Consequently, we computed pairwise correlations between the *K* predicted FCs leading to a *K* × *K* similarity matrix for each test subject. We then calculated the element-wise mean (8(a)) and standard deviation (8(b)) across the 23 similarity matrices. As shown in the figure, the dominant identity matrix pattern observed in the mean matrix combined with low values in the standard deviation matrix, verifies that the predictions are indeed distinct from one another.

**Figure 7:**
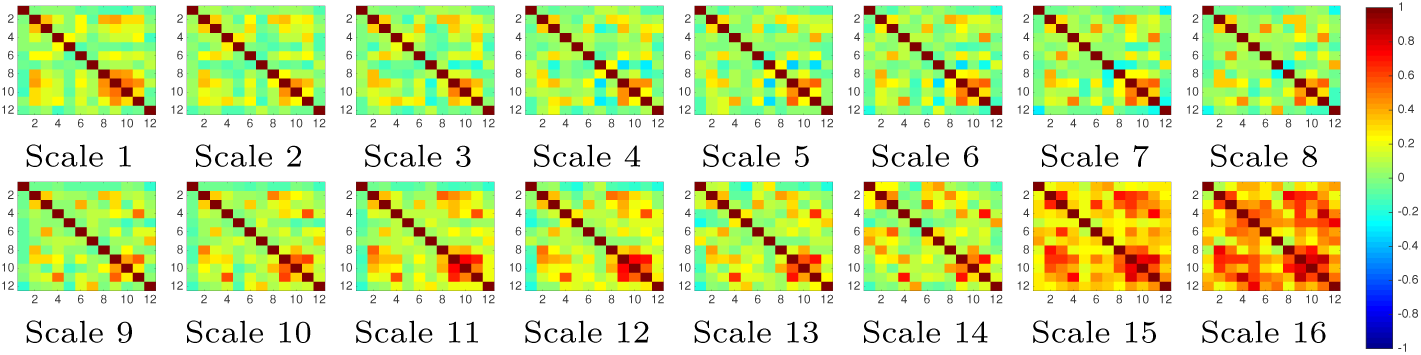
Distinctness of 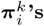. After state-specific MKL models are learned, we check the distinctness of 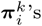 for every scale value ranging from *i* = 1,…, *m*(= 16) using Pearson correlation coefficient between every pair of states. Each of these m matrices is a *K*(*=* 12) × *K* similarity matrix. Distinctness of 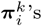 would ideally result in a *K × K* identity matrix and any deviation would indicate lack of distinctness of the learned 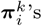. As observed in most of these *m* similarity matrices, majority of the off-diagonal entries in these pairwise correlation matrices are zero, indicating the distinctness 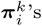. They are significantly distinct for global scales (scale indices *i* = 1,…, 8 in the top row) in comparison to local scales (scale indices *i* = 9,…, 16 in the bottom row), where they appear to be similar.

In the third experiment, we evaluated the accuracy of the predicted state-specific assignments of the proposed model. A wFC in the training phase is assigned to a state which it belongs to with the maximum probability of belongingness as described in 1 in section 2.2.2. The cluster assignments of wFCs to states should obey the principle of maximum intra-state similarity as well as maximum inter-state dissimilarity. Therefore the predicted FC for a state in the testing phase should have maximum similarity with the training wFCs belonging to the same state and also minimum similarity with training wFCs belonging to other states. For each predicted state-specific FC, a set of training wFCs is computed which lie in its proximity in the original space (of size *^n^*^(^*^n^*^−1)^/2). The mode of the wFC state-labels of this set of neighbouring training wFCs would indicate the estimated state label for the predicted FC. Recall from step 6. of Figure 1 that the tMKL model implicitly assigns a state-label to the predicted FCs. In this experiment, our aim is to compare the implicit label with the estimated label. The concurrence is measured through a confusion matrix aggregated over all the test subjects (see Figure 8(c)). The accuracy of the predicted state-specific assignment measures the number of instances for which this estimated state label matches the implicit state label. The confusion matrix (Figure 8(c)) has a dominant main diagonal and low off-diagonal elements, indicating that the implicit assignments seem to be valid. Overall accuracy of the state-specific assignments for the test subjects works out to be 87.68%.

**Figure 8:**
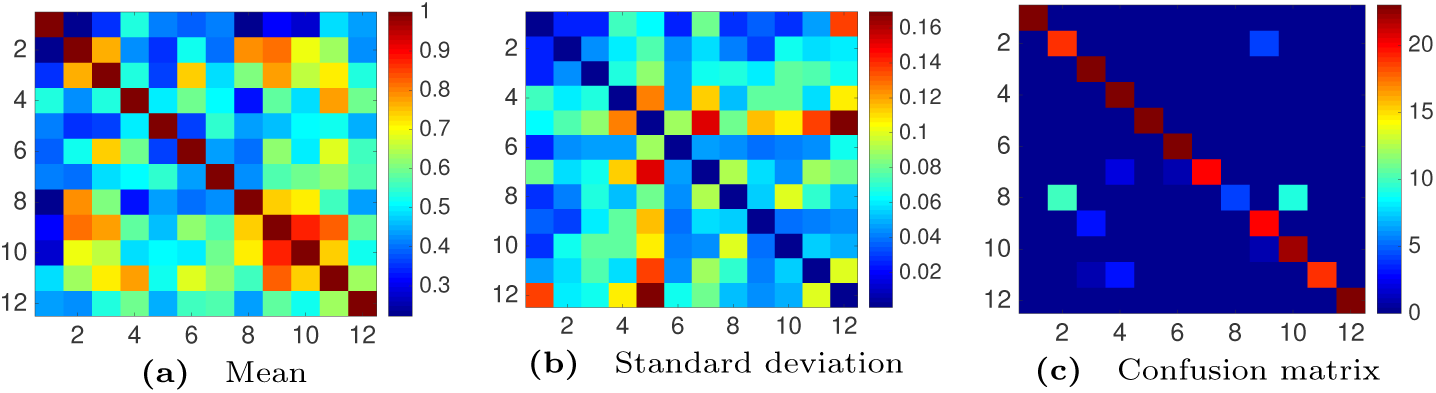
Quantitative state specificity of the model. Pearson correlation between all the possible pairs of the *K* state-specific FCs for a subject was calculated and stored in a *K × K* matrix. Element-wise mean and standard deviation across all the subject-specific matrices is shown in (a) and (b) respectively. The confusion matrix in (c) can be used for measuring the overall accuracy of state-specific predictions, and precision and recall for each state. For each testing subject there are *K* number of state-specific FCs. The state label against which a state-specific FC is predicted from tMKL model serves as the ground truth for this experiment. Empirically, each state-specific FC must be nearer to the training wFCs belonging to that state than from the training wFCs belonging to other states, thus attesting the accuracy of model prediction. As the manifold was constructed based on *L*1 similarity, we found the neighbors of the predicted FCs in the original space (of size *^n^*^(^*^n^*^−1)^/2). For this purpose, we searched for 25 nearest neighbors for each state-specific prediction and voted for the empirical state-belongingness. Rows (columns) in the confusion matrix depict the actual (predicted) states. Overall accuracy of tMKL model prediction for all the test subjects is 87.68%. It can be seen that non-zero off-diagonal entries result in reduced accuracy. To get a subject-specific measure of the state-specificity, we ran the same experiment for all the testing subjects independently. Noticeably, mean matrix is similar to the confusion matrix with very less standard deviation.

In summary, the above-mentioned experiments establish the ‘distinctness’ of states and their corresponding predictions.

## 4. Discussion

Besides understanding the relationship between the anatomical architecture and the functional dependencies, over the last decade, characterization of the temporal richness of the resting state functional MRI signal also has been a major trend in the field of cognitive neuroscience. Several approaches have been proposed to understand the inherent richness observed in the spontaneous spatio-temporal BOLD activity. Operator-based formulations of neural dynamics [11, 12] propose a generative model to predict functional connectivity from the structural connectivity via incorporating temporal dynamics into the model. Another class of techniques introducing spectral graph theoretic methods [16, 14, 18, 17] primarily focus on mapping the eigen-spectrum of SC and FC of individual subjects, but with minimal focus on the temporal richness. Here, we have proposed an innovative method which combines both anatomical constraints as well as incorporating temporal richness present in the endogenous activity. More specifically, our proposed model learns parameters specific to these latent states using a temporal Multiple Kernel Learning (tMKL) and finally predicts the grand average functional connectivity (FC) of the unseen subjects by employing a state transition Markov model. One of the interesting proposal in the framework is that tMKL learns a mapping between the underlying anatomical network and the temporal structure present in the empirical data to quantify gFC. Further, we have introduced a learning framework to find model specific parameters via state-specific optimization formulations and yet the model performs at par or better than state-of-the-art models for predicting the gFC. Moreover, our proposed model shows sensitivity towards individual subject’s SC as we have clearly demonstrated here with perturbation experiments. However, before we can clearly appreciate the novelty in the proposed techniques we need to understand the existing methods of relating underlying SC with windowed FC.

**Figure 9:**
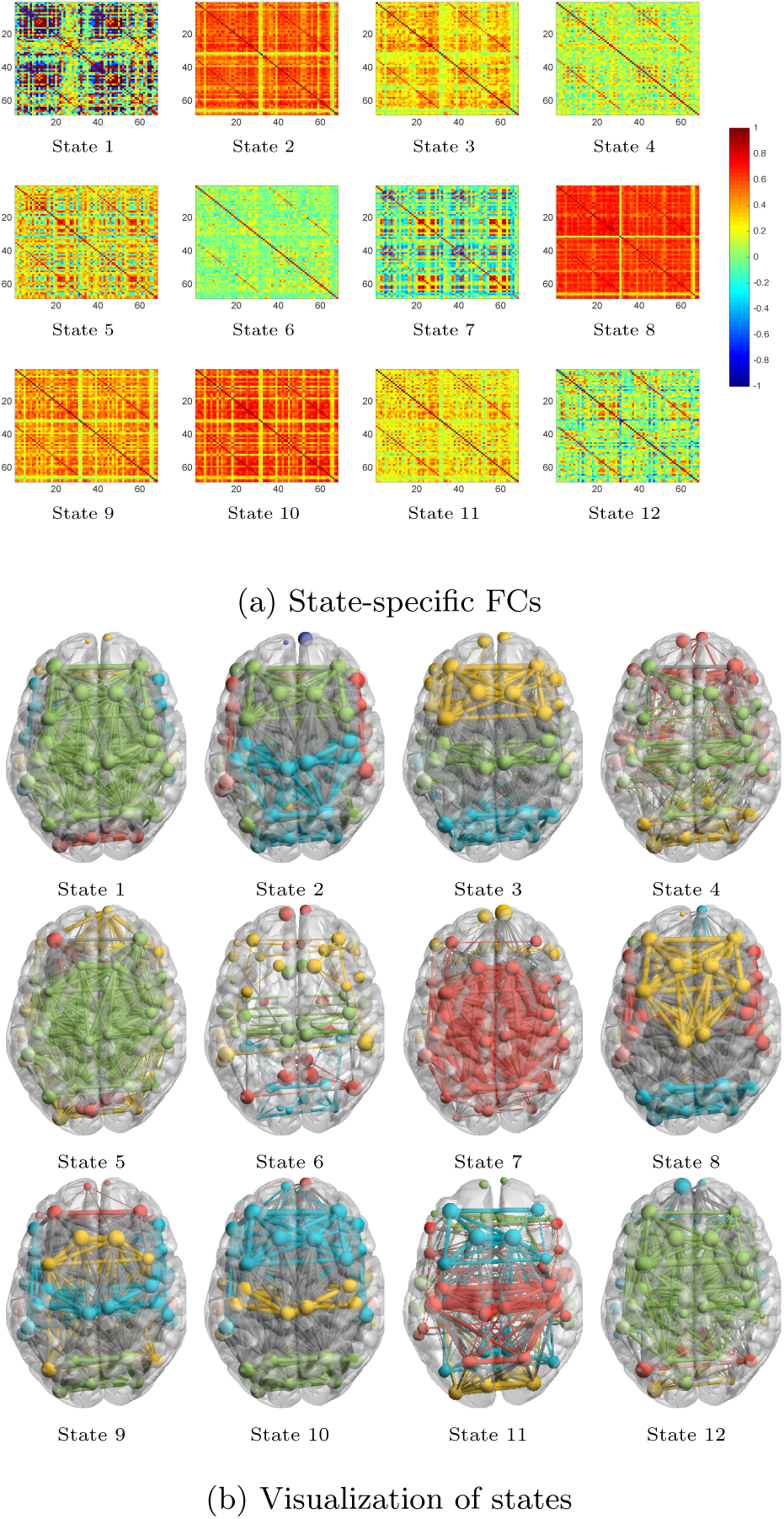
Qualitative state specificity. State specific FCs predicted for every subject are averaged across all testing subjects. (a) Visually distinct FC matrices are shown for all the 12 states. (b) Communities are identified for these mean FCs using Louvian algorithm available in brain-connectivity-toolbox [37] and Brain-net-viewer [38] was used for visualization of these communities. The distinct community structures clearly suggest that transient states are modeling different brain dynamics.

### 4.1. Relating underlying structural connectivity to Windowed FC

A different line of study by Allen et al. [21] and subsequent works by these and other authors have focused on the temporal structure of the windowed FCs (wFCs) and were able to successfully characterize state transitions, but without specifically relating the temporal dynamics to the underlying structure [39, 40, 41, 20]. Allen et al. collected all the wFCs from the training subjects and learned a k-means clustering model to discover distinct latent states (common to the cohort) that are visited by the brain. Our preliminary attempts at fusing SC using MKL model [17] with the temporal structure learned with k-means did not give satisfactory results (summarized in Figure S1 in Supplementary materials and the description therein). The proposed temporal multiple kernel learning (tMKL) model in this paper belongs to the class of spectral graph theoretic methods [16, 14, 18, 17]. The model attempts both at improving upon the quality of SC-FC mapping and also aims to characterize the temporal richness of the signal while incorporating the structural information in a principled way. The proposed temporal multiple kernel learning (tMKL) model is an attempt towards generating BOLD time series of a subject using only the SC. As summarized in Figure 1, the proposed pipeline partitions the BOLD time series of a subject into windows yielding wFCs. The underlying structure of the wFCs was learned via a manifold whose structure was further parameterized using GMM. The GMM components were hypothesized to be the states whose temporal evolution was succinctly captured in a Markov chain transition matrix. For each state, MKL model was learned to capture the SC-dFC relationship. The learned model was utilized to predict state-specific FCs for a test subject. These predicted FCs were further factored into latent time series and concatenated using the steady-state properties of the transition matrix. Pearson correlation of this final time series generates the predicted FC for a subject.

### 4.2. Rationale for t-MKL pipeline to discover latent temporal structure

In the following we will explain the rationale for various steps in the proposed pipeline. It appears that while modeling dFC using unsupervised techniques for clustering wFCs into states, one faces the curse of dimensionality problem head-on. During clustering, wFCs ought to be assigned to the same state as that of their neighbors because they are temporally contiguous and might share similarities. As we can see, wFCs lie in a high dimensional space, but based on their similarity with respect to their neighbors, they may lie on an intrinsic lower-dimensional manifold. This lower dimensional manifold becomes the space over which temporal structure could be precisely identified. Spectral embedding techniques utilize the similarity between the neighboring wFCs to discover the underlying manifold. After representing the temporal structure as a manifold, the next task is to parameterize the lower-dimensional structure. Once we obtain a lower-dimensional embedding, we need to cluster the wFCs to discover the discrete state space. Unsupervised approaches such as K-means clustering would yield spherical clusters, limiting the shape and size of states, whereas GMM clustering is a generalized clustering scheme. We parameterize the local density-distribution of wFCs over the manifold to a factor analysis model that further represents the manifold as a set of component Gaussians at various locations whose shape, orientation, and size depend on the local densities of the wFCs.

The proposed model is cohort-based and hence the underlying assumption is of the generalizability of the model to unseen test subjects. We have learned the Markov transition probability matrix on training wFCs and used this to generate long sequences of time series for test subjects eventually yielding a good approximation of the grand average FC with a maximum of 0.8 (see Figure 3).

### 4.3. Reproducibility of latent states and FC configurations

After presenting the rationale behind designing the proposed model that is shown to be successful at mapping SC-dFC-FC tripartite relationship, several expectations arise such as reproducibility of discovered states and their corresponding predicted FCs, sensitivity of the model to the underlying anatomical structure, state-specificity of the tMKL model and importantly verifying that the model does not overfit the training data. Several experiments were conducted in order to verify that the model satisfies these claims and the results presented in Section 3 point to the robustness of the performance of the model. Further, in order to verify whether the state-specific FCs predicted for a subject are distinct, we performed community detection over the mean state-specific FCs of the test cohort (see Figure 9). As can be clearly seen in the figure, regions in each state show distinct interaction patterns among themselves. The states seem to characterize the transient relationship among the ROIs which appear and disappear across the duration of the resting state scan. The markov chain state transition model further allows the characterization of the temporal fluctuations of the states that approximates latent temporal structure. Significantly, the MKL models were learned only over the individual states without any global error measure governing the learning process. Yet, the grand average FC prediction is at par or better than that of the MKL model and superior to the other competing approaches.

### 4.4. Conclusion

As part of future work, it will be interesting to explore the biophysical meaning of the model parameters. Other direction could be to characterize the dynamics better by predicting the time series itself rather than working with correlation matrices. An immediate investigation would be to explore the relationship between the latent time-series and the actual BOLD time-series. Another line of work would be to apply the proposed model to characterize dFC in various conditions such as neurodegenrative and psychiatric disease, healthy and pathological aging etc.

## 5. Acknowledgements

The authors would like to thank Arpan Banerjee for his insightful feedback and proofreading the manuscript. DR is supported by the Ramalingaswami Fellowship (BT/RLF/Re-entry/07/2014) from Department of Biotechnology (DBT), Ministry of Science & Technology, Government of India.

